# In vivo delivery of functional Cas:DNA nucleoprotein complexes into recipient bacteria through a Type IV Secretion System

**DOI:** 10.1101/2024.01.18.576218

**Authors:** Dolores L. Guzmán-Herrador, Andrea Fernández-Gómez, Florence Depardieu, David Bikard, Matxalen Llosa

## Abstract

CRISPR-associated (Cas) endonucleases and their derivatives are widespread tools for the targeted genetic modification of both prokaryotic and eukaryotic genomes. A critical step of all CRISPR-Cas technologies is the delivery of the Cas endonuclease to the target cell. Here, we investigate the possibility of using bacterial conjugation to translocate Cas proteins into recipient bacteria. Conjugative relaxases are translocated through a Type IV Secretion System (T4SS) into the recipient cell, covalently attached to the transferred DNA strand. We fused relaxase R388-TrwC with the class 2 Cas endonuclease Cas12a and confirmed that it can be transported through a T4SS. The fusion protein maintained its activity upon translocation by conjugation into the recipient cell, as evidenced by the induction of the SOS signal resulting from the cuts produced by the endonuclease in the recipient cell, and the detection of mutations at the target position. We further show how a template DNA provided on the transferred DNA can be used to introduce specific mutations. The gRNA can also be encoded by the transferred DNA, enabling its production in the recipient cells where it can form a complex with the Cas nuclease transferred as a protein. This self-contained setup enables to target wild type bacterial cells. Finally, we extended this strategy to the delivery of relaxases fused to base editors. Using both TrwC and MobA relaxases as drivers, we achieved precise editing of transconjugants. Thus, conjugation provides an *in vivo* delivery system for Cas-derived editing tools, bypassing the need to deliver and express a *cas* gene in the target cells.

**Significance Statement:** We have developed a novel approach for introducing CRISPR-Cas genetic tools into bacteria. During bacterial conjugation, the relaxase protein is transferred through the secretion system covalently attached to the transferred DNA. By fusing the Cas protein with the conjugative relaxase, we have observed functional Cas activity in the recipient cells, eliminating the need for nuclease expression in these cells. The covalently attached DNA molecule can supply gRNA and donor DNA, enabling seamless genetic modifications through recombination. We have also successfully translocated fusions of relaxases to base editors which are active in recipient cells. This method can be applied to any potential recipient cells, making it particularly interesting for wild type bacterial strains that lack available genetic tools. Furthermore, this method has the potential to be extended to eukaryotic cells.

## Introduction

The field of genetic engineering has been significantly transformed by the advent of CRISPR-Cas systems, which have provided a powerful and precise tool for genome manipulation. These systems have not only revolutionized the landscape of biotechnology but have also given rise to a new era of genomic research. Biotechnological applications mostly rely on class 2 systems, where a single Cas protein is sufficient to perform specific cleavage of target nucleic acids (1). Additionally, dead versions of these endonucleases can be used to deliver fused peptides to specific genomic locations, leading to site-specific actions such as activation/repression of transcription or histone modification, depending on the protein fused to the Cas protein (2–4). One such modification involves the utilization of base editors (BE), arising from the fusion of engineered Cas proteins like nickase Cas (nCas) or non-catalytic Cas (dead Cas, dCas), with a deaminase enzyme, either cytidine or adenine deaminases. This results in cytidine (CBE) or adenine base editors (ABE), respectively. These innovative systems facilitate the targeted alteration of specific nucleotides, enabling the conversion of cytidine to thymidine via CBE or adenine to guanine via ABE at precise genomic positions (5). BE have been proved to be more efficient compared to the canonical CRISPR-Cas system in gene editing (6, 7).

The development of CRISPR-Cas tools and therapies relies on the identification of delivery strategies that are both efficient and safe (reviewed by (8–12)). The most widely used strategy involves the introduction of DNA to express the Cas protein and gRNA in target cells, which requires controlled and tunable expression of the *cas* gene. However, this can be challenging in non-model bacteria, which often have restriction-modification systems that degrade incoming DNA. Moreover, replicative plasmids and selectable markers are frequently not available, and commonly used inducible expression systems may not be functional, while the endogenous regulatory sequences can be difficult to identify (13). An excess of *cas* expression over time may lead to increased off-target effects and toxicity (14). The alternative approach involves introducing the Cas protein and gRNA either as mRNA/gRNA or as a ribonucleoprotein. While a variety of strategies have been developed for this latter technique, achieving effective intracellular delivery, especially in vivo, is still challenging (15). Ideally, a new system that could produce and deliver the Cas protein in vivo would bypass these problems. With this goal in mind, we have tested the possibility of sending Cas proteins fused to conjugative relaxases, proteins that are naturally translocated between bacterial cells during conjugation.

Bacterial conjugation is an efficient method of introducing DNA into bacteria, including strains and species that can be difficult to transform (16, 17). Donor and recipient bacteria come into physical contact through a Type IV Secretion System (T4SS), a multiprotein complex that spans the inner and outer membrane of bacterial cells (18). A conjugative relaxase recognizes and cleaves its target (the *oriT* sequence) in the DNA strand to be transferred, making a covalent bond with its 5’end. This nucleoprotein complex is recruited by the T4SS and translocated to the recipient cell, where the relaxase catalyzes the recircularization of the transferred DNA strand (19). Some broad host range conjugative systems, such as that of plasmid RP4, have been used to deliver CRISPR-Cas systems in a large number of genetically amenable bacteria (20–23). We reasoned that we could deliver the Cas protein by conjugation fusing it to the conjugative relaxase, which is translocated through the T4SS. There is a precedent of Cas translocation through a T4SS: A previous work used the VirB/D4 T4SS of *Agrobacterium tumefaciens* to translocate the Cas9 protein fused to the translocation peptide signal VirF into eukaryotic cells (24). In this work, we generate fusion proteins between conjugative relaxases and Cas12a endonuclease, and we show that the translocated protein can cleave and introduce mutations at a target position. Moreover, we also generate fusions between relaxases and base editors, resulting in the precise editing of the target locus in transconjugants. Additionally, we show that the covalently attached transferred DNA strand can be used to encode a gRNA and a template DNA to introduce specific mutations in the target gene of a wild-type recipient bacterium. Thus, we have generated an *in vivo* method of delivering Cas-derived genetic tools that bypasses the need to express the *cas* gene in the target cell.

## Results

### Construction and validation of a TrwC-Cas12a fusion protein

We engineered a gene that encodes the AsCas12a protein with a C-terminal 3xHAtag fused to the C-terminus of the relaxase TrwC of the conjugative plasmid R388, resulting in the fusion protein TwC-Cas12a. This gene was cloned under the regulation of a P*tet* promoter into plasmid pTrwC-Cas12a (**Table 1**). The stability of the fusion protein was analysed by Western Blot. The results (**SI Appendix Fig. S1**) showed a band with the expected size for the fusion protein (263 kDa), as well as additional bands that likely correspond to degradation products or partially translated proteins. This confirms that while not fully stable, the full-length protein is produced, enabling the transfer of TrwC-Cas12a through the T4SS.

**Table 1.** Main plasmids used in this work*.

| Name in manuscript | Laboratory name | Main characteristic |
| --- | --- | --- |
| pgRNA-lacZ | pLG15 | *Plac lacZ<sub>gRNA</sub>* |
| pgRNA-sacB | pLG19 | *Plac sacB<sub>gRNA</sub>* |
| pTrwC-Cas12a | pLG24 | *Ptet trwC-cas12a* |
| pTrwC | pLG22 | *Ptac trwC* |
| pCas12a | pLG14 | *Ptac cas12a* |
| pSOS | pZA31 | *Psos gfp* |
| pgRNA-sacB-oriT | pLG29 | *oriT Plac sacB<sub>gRNA</sub>* |
| poriT | pSU1186 | *oriT* |
| pHR_oriT | pLG27 | *oriTR388+oriVR6K Ptac::sacB\**<br>homologous recombination<br>cassette |
| pBE | pLG32 | *Ptet dcpf1-be* |
| pMobA | pAF21 | *Ptet mobA* |
| pTrwC-BE | pLG33 | *Ptet trwC-dcpf1-be* |
| pMobA-BE | pAF18 | *Ptet mobA-dcpf1-be* |
| pgRNA-lacZ<sub>BE</sub> | pLG38 | *Plac lacZ<sub>gRNA-BE</sub>* |
| pgRNA-ErmB<sub>BE</sub> | pAF23 | *Plac ermB<sub>gRNA-BE</sub>* |
| pErmB | pAF20-ATG | *Pbad ermB* |
| pErmB* | pAF20 | *Pbad ermB with no ATG\** |
| RSF1010mobA- | pMTX808 | *pAA58::ApR in mobA* |
| R388trwC- | pSU1445 | *R388::Tn5tac1 in trwC* |
| pUC8 | pUC8 | Multicopy ApR cloning vector |

\*Detailed description of the plasmids in *SI Appendix Table S3*

The functionality of TrwC in the fusion protein was tested by complementation assays of a R388 *trwC* deficient mutant. The results (**SI Appendix, Table S1**) showed that the conjugation frequencies were similar to those obtained with TrwC. We then tested if TrwC-Cas12a maintains RNA-guided DNA cleavage activity by a lethality assay, since the introduction of double strand breaks by Cas proteins in the chromosome of *E. coli* leads to cell death (25). We co-electroporated into *Escherichia coli* D1210 plasmids pTrwC-Cas12a and either pgRNA-lacZ or pgRNA-sacB, which encode a gRNA under the control of the IPTG-inducible promoter P*lac*, targeting a chromosomal gene (*lacZ)* or a gene not present in the chromosome (*sacB),* respectively. The resulting transformants were selected on plates with or without induction. No transformants were obtained in the presence of the target and upon induction of the fusion protein (**SI Appendix, Fig. S1**), confirming that TrwC-Cas12a can cleave target genomic sequences when an appropriate gRNA is present in the cell, and that its expression is tightly regulated under the control of the P*tet* promoter.

### Endonuclease activity of TrwC-Cas12a in the recipient cell

Observing TrwC-Cas12a activity after translocation into the recipient cell presents a greater challenge, as only a single or few TrwC-Cas12a proteins are expected to be transferred via conjugation, as opposed to the continuous protein production upon electroporation. To test if TrwC-Cas12a could be translocated through the T4SS of R388 during conjugation and recover its activity in the recipient, we used two different strategies, as depicted in **Fig. 1**. Initially, we measured the induction of the SOS response in transconjugants, which is expected to be triggered after the introduction of DSB by Cas12a (25). Subsequently, we isolated mutants that inactivate the target gene and characterized them. Cas12a cleavage should result in a specific mutational signature at the target position.

**Figure 1.**
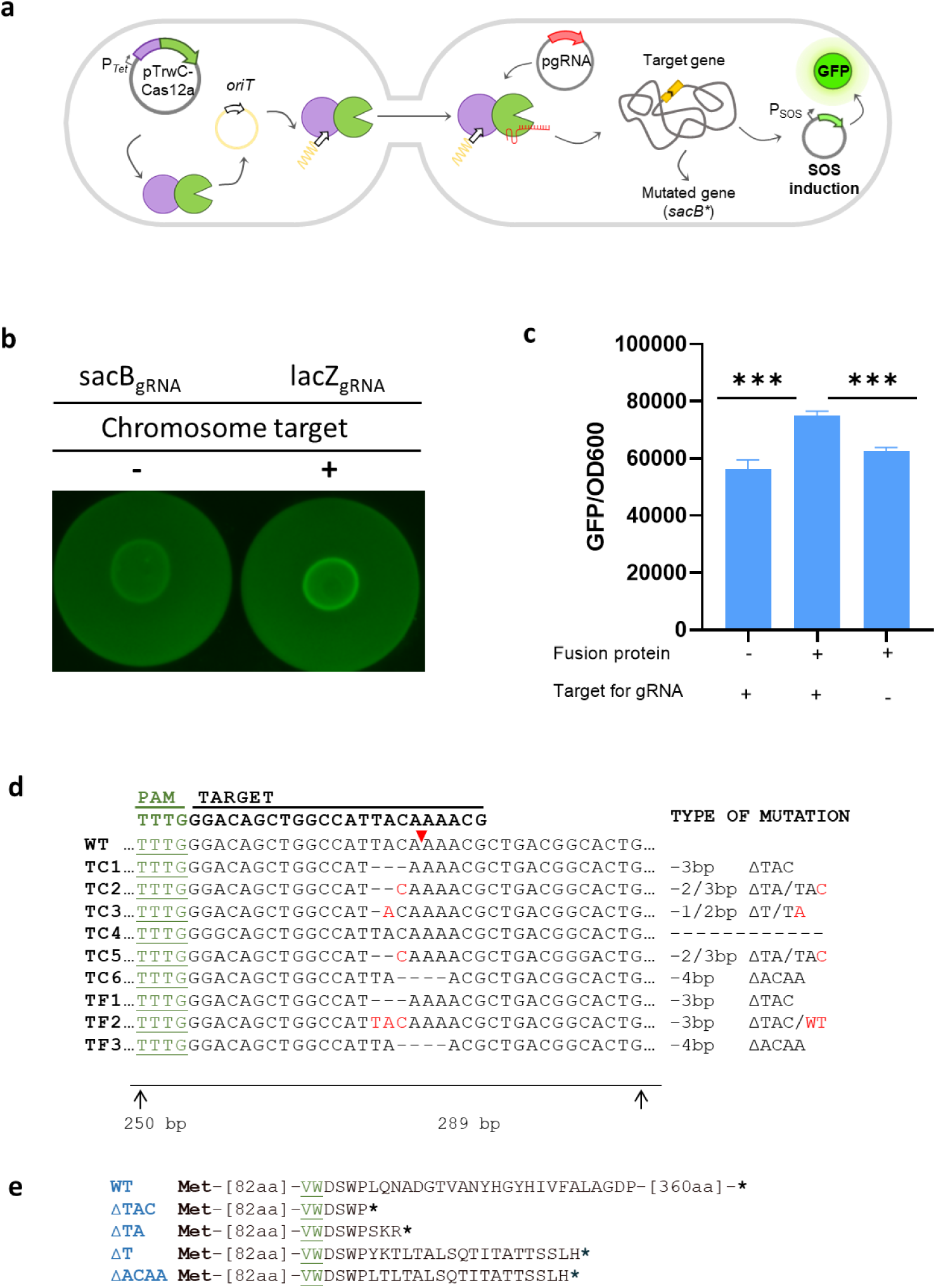
Endonuclease activity of TrwC-Cas12a in the recipient cell. **a)** Schematic representation of Cas12a activity assays in the recipient cell. In the donor cell, pTrwC-Cas12a (in grey) will express TrwC-Cas12a. Thanks to its relaxase activity, the fusion protein will cleave and bind covalently to the *oriT* (white arrow), and the complex will be recruited and translocated through the T4SS into the recipient cell. In the recipient, pgRNA-lacZ or pgRNA-sacB (in red) will produce a gRNA targeting a gene on the chromosome. Thanks to its endonuclease activity, the incoming TrwC-Cas12a will process the gRNA generating a complex, which will be guided to the target gene, where it will produce a DSB. **b)** Induction of SOS response by TrwC-Cas12a in the recipient cell. Visualization of conjugation filters under a fluorescent lector system after 3h of mating under induction conditions. Filter on the right (gRNA with a target on the chromosome) shows increased fluorescence level in comparison with the filter on the left (gRNA without target on the chromosome). **c)** GFP fluorescence relative to OD_600_ levels, measured with a TECAN Infinite M200 Pro. Data correspond to the mean of 3 independent assays (***, P < 0.0005; *, P < 0.05). **d)** Analysis of the *sacB* mutations in sucrose-resistant colonies. Alignment of the *sacB* region close to the PAM and target sites for Cas12a-gRNA. The *sacB* region was PCR-amplified from sucrose-resistant transconjugants (TC1 – TC6) and from sucrose-resistant transformants (TF1-TF3). The *sacB* sequence in strain MG1655::*sacB* was also determined and is shown at the top for comparison (WT). The PAM sequence and the spacer sequence are shown at the top. The red triangle marks Cas12a cleavage site in the shown DNA strand. Nucleotides in red mark the site where the DNA sequence splits into two (see text). **e**) Amino acid sequences of SacB variants resulting from the different mutations. The deletions are indicated at the left, in blue. The amino acids shown in green and underlined are encoded by the PAM sequence. Stop codons are shown as **\***.

We conducted conjugation assays using as donor *E. coli* D1210 harbouring R388*trwC-* complemented with pTrwC-Cas12a and as recipient MG1655 harbouring a reporter plasmid with a *gfp* gene under the control of a SOS-responsive promoter (pSOS), and the pgRNA. In order to discard an increase of the SOS signal triggered by the conjugation process itself (26), we also performed mating using D1210 harbouring R388*trwC-* without plasmid complementation. After the matings, the level of GFP was visualized in recipients that express either pgRNA-sacB (without chromosomal target) or pgRNA-lacZ (with chromosomal target) (**Fig. 1b**). We detected a significant increase in fluorescence when TrwC-Cas12a was translocated into recipients expressing the gRNA against *lacZ* (**Fig. 1c**). These data demonstrate that Cas12a induces the SOS response and that TrwC-Cas12a is active as a site-specific endonuclease in the recipient cell after translocation through the T4SS.

To obtain direct evidence for Cas12a mutagenic activity in the recipient cell, we aimed to select mutations introduced as a result of DNA repair following Cas12a cleavage at the target position. Sequence rearrangements have been reported following Cas9 cleavage in the chromosome of *E. coli* (25). To investigate if such rearrangements could occur after cleavage by TrwC-Cas12a, we selected transconjugants resistant to sucrose in a *sacB*-containing strain. The expression of this gene in the presence of sucrose is lethal in bacteria (27), so the transconjugants will only survive if there is a mutation inactivating the gene. We conducted several matings translocating the fusion protein or the relaxase into target or non-target recipients (MG1655::*sacB* and MG1655, respectively) and no differences in conjugation frequencies were observed among the different conditions (not shown). However, when we selected for sucrose-resistant transconjugants, several colonies appeared when TrwC-Cas12a was translocated into the recipient containing *sacB*. After confirming by PCR amplification that the *cas12a* gene was not present in the transconjugants, 11 sucrose-resistant transconjugants were analyzed by PCR amplification of the 5’ end of the *sacB* gene. In 5 cases, we did not observe any visible PCR product, possibly due to deletions encompassing the region amplified with the primers. The sequence of the PCR products of the 6 remaining transconjugants was determined (**Fig. 1d**). All transconjugants but one showed 1-4 nt deletions in the target DNA sequence. Interestingly, the sequences of TC2, TC3 and TC5 showed a mixture of two DNA sequences after the Cas12a cleavage site, which corresponded to two different deletions (shown in red in the **Fig. 1d**). These mixed colonies likely indicate the generation of different mutations on different copies of the chromosome in the recipient cell. All the deletions produced an early stop codon on the *sacB* sequence (**Fig. 1e**). To discard a role of TrwC in the cleavage pattern, we compared the *sacB* mutations of the transconjugants with the ones produced by the cleavage of Cas12a alone. We co-electroporated pCas12a and pgRNA-sacB into MG1655::*sacB,* and plated the transformants in sucrose-containing medium. Three sucrose survival transformants were selected. We amplified the *sacB* region, and determined the DNA sequence (**Fig. 1d**, TF1-TF3). We detected 3-nt and 4-nt deletions at the target sequence, which coincided with the ones observed in several transconjugants. Thus, we confirm that neither conjugative DNA transfer nor TrwC activity is involved in the type of mutations obtained, which are solely derived from Cas12a endonuclease activity. In summary, these results provide direct proof of the Cas12a activity of TrwC-Cas12a in the recipient cell after translocation through the T4SS.

### Use of donor DNA molecule to encode the gRNA and to provide template DNA

Since the relaxase is translocated covalently attached to a DNA strand during conjugation, we used this DNA as a source for the other elements of the genetic modification machinery: the gRNA or a template DNA to promote homologous recombination-mediated gene editing. This method allows us to target unmodified recipient cells, since we send all the components from the donor bacteria (**Fig. 2a**).

**Figure 2.**
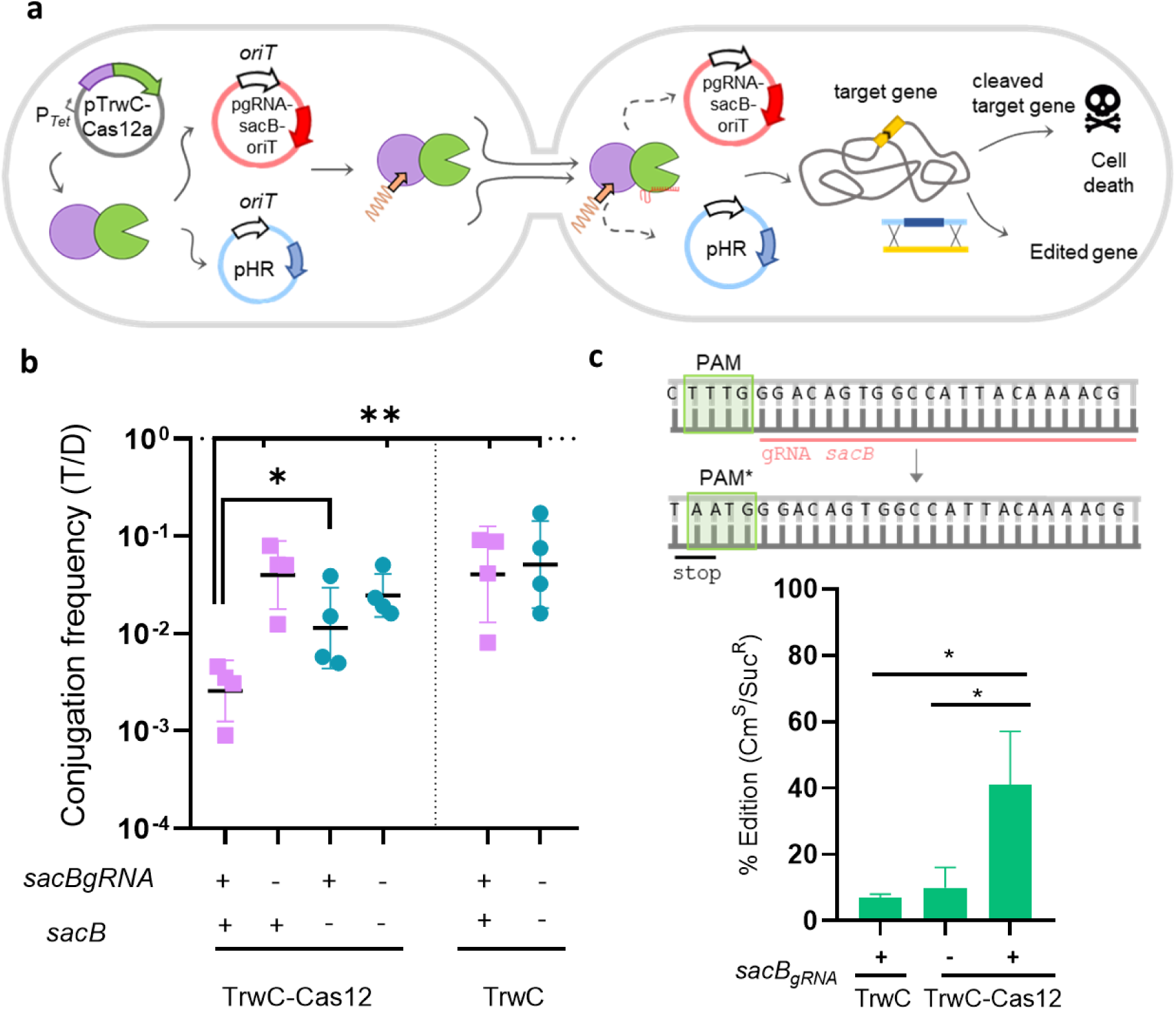
Translocation of donor DNA encoding the gRNA or a DNA template. **a)** Schematic representation of the assays. In the donor cell, pTrwC-Cas12a (in grey) will express TrwC-Cas12a. Thanks to its relaxase activity, the fusion protein will cleave and bind covalently to the pgRNA-sacB-oriT plasmid (in red) or to the pHR_oriT (in blue), and the complex will be recruited. The complex formed by the fusion protein and the mobilizable ssDNA of interest (represented as squiggly orange line) will be translocated through the T4SS into the recipient cell. In the recipient cell, if the attached DNA is the pgRNA-sacB-oriT, the incoming complex will be guided to the target gene, where it will produce a DSB, which is lethal to the bacteria. On the other hand, if the translocated plasmid is the pHR_oriT, the DSB produced in the chromosome of the recipient bacteria will be repaired through homologous recombination, utilizing the HR cassette provided by pHR_oriT, generating specific editions in this gene. **b)** Lethality in transconjugants upon mobilization of a plasmid encoding the gRNA. T/D, transconjugants/donor. Data correspond to 4 independent assays (**, P < 0.005; *, P < 0.05). **c)** Introduction of seamless mutations using a HR cassette. *Top*, the genome sequence of MG1655::*sacB* and the design of the *sacB* homologous recombination (HR) cassette, where the PAM sequence was mutated to generate an early STOP codon (PAM*). *Bottom*, percentage of edition with TrwC or TrwC-Cas12a.The editing rate is calculated as the fraction of chloramphenicol sensitive colonies which had incorporated the *sacB* mutations from the HR cassette per sucrose resistant colony analyzed. *sacB*_gRNA_ presence in the recipient cell is indicated with a + or a -. The data represent the results of three independent experiments (*, P < 0.05).

We constructed a mobilizable plasmid encoding the gRNA against the *sacB* gene (pgRNA-sacB-oriT). This plasmid is translocated covalently attached to TrwC-Cas12a into the recipient cell, where the gRNA will be expressed. We generated donor bacteria carrying 3 plasmids: R388*trwC*, pTrwC-Cas12a or pTrwC (negative control for Cas12a activity), and pgRNA-sacB-oriT or poriT (a mobilizable plasmid that does not encode a gRNA for Cas12a). As recipient strains, we used MG1655::*sacB* or MG1655, with and without a target gene, respectively. The results (**Fig. 2b**) showed a significant decrease in the number of transconjugants only when the gRNA was translocated by TrwC-Cas12a into a recipient cell containing the target gene (*sacB*), indicating that the endonuclease is introducing the DSB in its target.

The CRISPR-Cas gene editing system can edit a cell without leaving a scar if provided with a homologous template carrying the desired mutation. To test if the DNA template could be provided in the donor DNA, we constructed plasmid pHR_oriT. This is a mobilizable suicide plasmid that contains a *sacB* homologous recombination cassette under the control of a Tac promoter and with the PAM sequence mutated to generate an early STOP codon in the gene (**SI Appendix Fig. S2a**). Mobilization of the plasmid using TrwC or TrwC-Cas12a was equally efficient (**SI Appendix Table S1**). If TrwC-Cas12a promotes recombination, we should detect an increase in the editing efficiency in comparison with the spontaneous recombination of the template by homologous recombination. We performed matings using DH5αpir as donors, mobilizing pHR_oriT with TrwC or TrwC-Cas12a. As recipient cells, we used MG1655*::sacB* harboring either pgRNA-sacB or pUC8. **Table 2** summarizes the results of 3 independent assays. We did not observe differences in the frequency of sucrose-resistant recipients when TrwC-Cas12a was delivered to the cell compared to TrwC alone (**Table 2**, Sucrose^R^ frequency column). However, when we analyzed the colonies for edition, we found a substantial increase in the fraction of sucrose resistant colonies that had incorporated the desired mutation. Cm-sensitive colonies were selected to discard integrants (as schematized in **SI Appendix Fig. S2b**), and the region containing PAM was amplified to determine if the colonies had incorporated the PAM mutation from the recombination cassette. **Fig. 2c** shows the editing ratio, calculated as the number of colonies which had incorporated the desired mutation, divided by the total number of Sucr^R^ colonies analysed for that condition. We could see a ca. 5-fold increase in the editing ratio when both TrwC-Cas12a and pgRNA-sacB were present. As expected, some edited colonies could also be detected when TrwC and the pgRNA-sacB were delivered due to the homologous recombination background, which can occur in the absence of DNA cleavage. Overall, these results demonstrate that TrwC-Cas12a promotes recombination of the HR template that was co-delivered with it in the recipient cell in the presence of the specific gRNA.

**Table 2.** Homologous recombination assays.

| Relaxase<sup>1</sup> | Recipient<sup>2</sup> | Sucrose<sup>R</sup><br>frequency<sup>3</sup> | Cm<sup>S</sup>/Suc<sup>R</sup><sup>4</sup> | %Edition<sup>5</sup> |
| --- | --- | --- | --- | --- |
| **TrwC** | *sacB<sub>gRNA</sub>* | $3.8 \times 10^{-5} \pm 3.5 \times 10^{-5}$ | 23/42 | 7.1 |
| **TrwC-Cas12a** | *sacB<sub>gRNA</sub>* | $1.7 \times 10^{-5} \pm 1.3 \times 10^{-5}$ | 34/42 | 40.5 |
| | no gRNA | $9.7 \times 10^{-6} \pm 4.2 \times 10^{-6}$ | 35/42 | 9.5 |

The data represent the results of four to six experiments.
<sup>1</sup> Donor cells were DH5αpir harboring pHR\_oriT and R388*trwC*- complemented with pTrwC or pTrwC-Cas12a.
<sup>2</sup> MG1655::*sacB* was used as recipient cell, harbouring pUC8 (no gRNA) or pgRNA-*sacB* (*sacB<sub>gRNA</sub>*).
<sup>3</sup> Expressed as sucrose-resistant recipients per recipient.
<sup>4</sup> Cm-sensitive per sucrose-resistant colonies
<sup>5</sup> Percentage of Cm<sup>S</sup> edited colonies per sucrose-resistant colonies

### Translocation of dCas12a fused to Base editors

In order to extend the Relaxase-Cas approach to other conjugative relaxases and other Cas-related genetic editing tools, our subsequent goal was to employ our delivery system to deliver a CBE. For this purpose, we used the dCpf1-BE system (hereafter referred to as BE), which comprises the cytidine deaminase APOBEC1, dLbCas12a and a Uracil Glycosylase Inhibitor (UGI) (28). We fused this BE to either TrwC or to the MobA relaxase from plasmid RSF1010 (plasmids pTrwC-BE and pMobA-BE, **Table 1**). The latter was chosen due to its smaller size, promiscuity and ability to be fused to translocation signals while retaining its functionality (29–31). Following the same approach used for the TrwC-Cas12a fusion, we inserted the BE gene at the 3’ end of the relaxase genes, eliminating the stop codon, downstream of the P*tet* promoter. We then verified the expression and stability of the newly formed TrwC-BE fusions (**Fig. S3**). As controls, we also cloned the BE and MobA separately under the regulation of the same promoter (pBE and pMobA, **Table 1**).

To test the relaxase activity in the fusion proteins, mating assays were conducted. TrwC and TrwC-BE complementation of R388*trwC-* plasmid, and MobA and MobA-BE complementation of RSF1010*mobA*-were tested in a strain providing the RP4 T4SS from the chromosome. Results demonstrated that both relaxase-CBE fusions exhibited the ability to mobilize a plasmid through complementation at a frequency comparable to that achieved with TrwC or MobA alone (**Table S1**).

Subsequently, we proceeded to test the functionality of dCpf1-BE, which had not been previously documented in bacteria. To achieve this, we engineered the pErmB* plasmid, incorporating a target that would allow us to identify a ‘gain-of-function’ mutation. The ATG start codon of the erythromycin (Em) resistance gene (*ermB*), under the regulation of an arabinose-inducible promoter, had been switched to ACG. Plasmid pgRNA-ermB_BE_ encoded a gRNA encompassing this mutation, so that edition would restore the ATG codon by changing C to T. Therefore, edited cells would be Em-resistant (**Fig. 3a**). We first assessed the BE activity of the constructs in *E. coli* by electroporation of the components of the system. Plasmids encoding either BE, TrwC-BE or MobA-BE, and MobA or TrwC as negative controls, were electroporated into DH5α cells containing plasmids pgRNA-ErmB_BE_ and the target pErmB*. These cells were then selected with Em and appropriate inducers. Following a 2-day incubation period, colonies appeared on plates containing the BE, either independently or as a fusion protein. The *ermB* gene was PCR-amplified from Em-resistant colonies, and DNA sequences showed that 90 to 100% had edited pErmB as expected, while no edition was observed in the occasional Em resistant colonies that emerge in the negative controls with no BE (**Fig. 3b**). We calculated the editing frequency per transformed cell as the ratio between Em-resistant and total transformants obtained (**Fig. 3c**), achieving a frequency of almost 100%.

**Figure 3.**
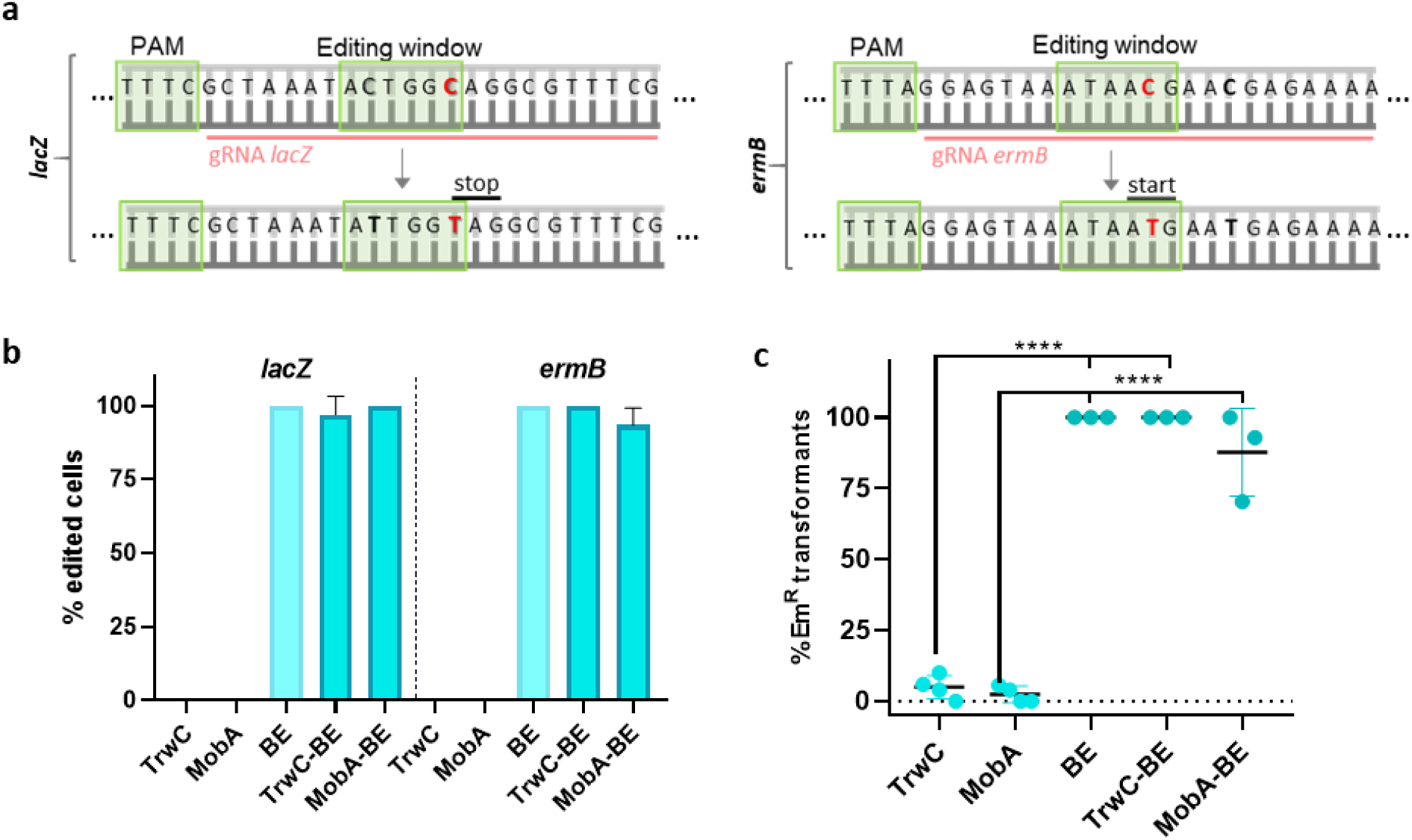
Activity of relaxase-BE fusions in *E. coli*. **a)** Sequence of D1210 *lacZ* gene (left) and mutated *ermB* gene present in plasmid pErmB* (right). The sequences complementary to the gRNAs against *lacZ* and *ermB* are indicated (red line). The PAM sequences and the editing windows are also highlighted (green boxes). The nucleotides where editing is expected to occur are marked in red, as well as the resulting stop or start codons. Other C nucleotides where edition was detected are highlighted in bold. **b)** Percentage of edition of selected colonies. Plasmids encoding BE, Relaxase-BE fusions and relaxases (as negative controls) were coelectroporated with a pgRNA plasmid into cells with the corresponding target gene (genomic *lacZ* or plasmidic *ermB\**). Selected colonies (white for *lacZ* and Em-resistant for *ermB\**) were sequenced to confirm the edition. Data correspond to the mean of at least 3 independent assays. Error bars represent SD. **c)** Percentage of Em-resistant transformants. Mean ± SD is represented. ****, P < 0.0001.

We also tested the system by editing the genomic *lacZ* gene. In this case, plasmid pgRNA-lacZ_BE_ encoded a gRNA with a C within the editing window which, upon changing to T, would result in a premature stop codon (**Fig. 3a**). It was cotransformed with BE, Relaxase-BE or relaxase plasmids into the *lacZ*-bearing strain D1210, and selected in X-Gal, so that successfully edited colonies would be white. We were able to detect lighter-shaded colonies in the cases where the BE or relaxase-BE were present, and sequencing of the *lacZ* region showed that 87.5 to 100% of those colonies were edited (**Fig. 3b**). Both BE and relaxase-BE fusions exhibited a preference for editing a C within the editing window which did not give rise to a stop codon. Nevertheless, the resultant colonies displayed a lighter phenotype, possibly attributable to a CRISPR interference phenomenon (29). Interestingly, when we extended the incubation of the plates for 3 days, a majority of the colonies eventually developed a lighter halo, indicating that edition was still ongoing after plating. For both *ermB* and *lacZ* edition, edited colonies showed a mixture of edited and unedited sequences. However, the edited one predominated after successive passages. Such mixed sequences have previously been reported in BE-induced mutations (30–32). Altogether, these findings demonstrate that the BE and its fusions to TrwC and MobA relaxases are active in *E. coli*, targeting both genomic and plasmidic genes.

Next, we aimed to assess the editing activity of the fusion proteins upon translocation into the recipient bacteria containing a gRNA targeting a chromosomal *lacZ* or plasmidic *ermB* gene, as depicted in **Fig. 4a**. For the matings targeting *lacZ*, donor bacteria harbored the R388*trwC-* plasmid, complemented by non-mobilizable plasmids pTrwC or pTrwC-BE. Selection of the transconjugants was done in X-Gal containing media (**Fig. 4b**). We sequenced the target position in 10 lighter-shaded transconjugants and detected modifications in 52% of them (**Fig. 4c**). In order to test the MobA-BE fusion against the plasmidic target *ermB*, donor cells had the RP4 system integrated into their chromosome plus mobilizable plasmid RSF1010*mobA*-, complemented with MobA or MobA-BE. Transconjugants were selected in the presence of Em, and after 2 days of incubation post-plating, Em-resistant colonies appeared in matings with MobA-BE but not MobA. DNA sequencing of these transconjugants revealed that, on average, 89% of them were edited (**Fig. 4c)**. To estimate the editing frequency per transconjugant, we calculated the ratio between Em-resistant and total transconjugants (**Fig. 4d**). The results revealed that, on average, 0.1% of the total transconjugants were edited.

**Figure 4.**
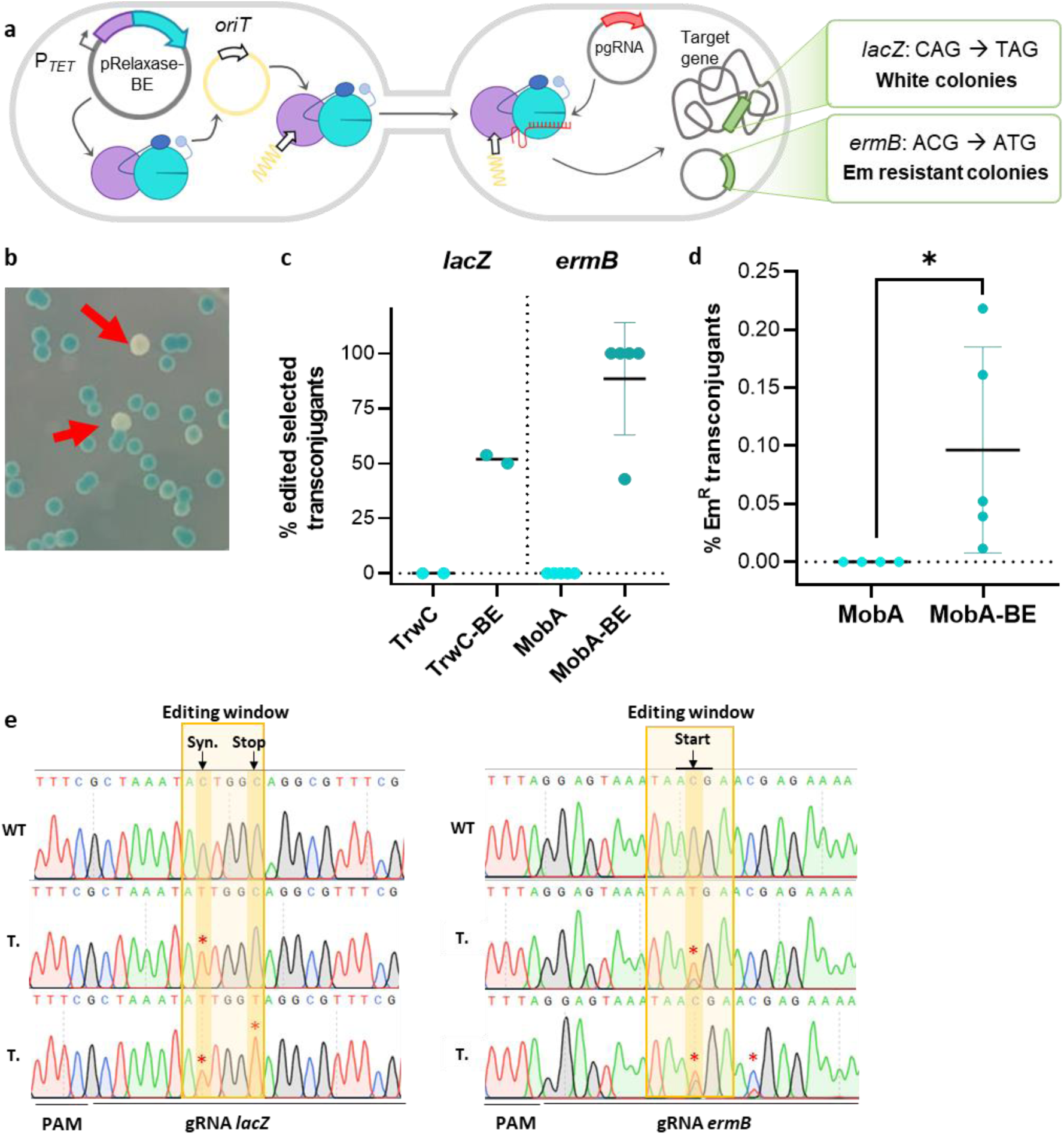
Activity of Relaxase-BE fusions delivered by conjugation. **a)** Schematic representation of the conjugative delivery and activity of Relaxase-BE fusions in the recipient cell. In the donor cell, pTrwC-BE or pMobA-BE express the Relaxase-BE fusions, which are then translocated to the recipient cell. In the recipient cell, the fusion protein binds to a gRNA produced by a pgRNA plasmid and targets a specific gene, where it performs base conversion. **b)** Transconjugants obtained by the translocation of TrwC-BE and selected in the presence of X-Gal. Red arrows indicate two light-colored colonies that were found to be edited. **c)** Editing efficiency of Relaxase-BE fusions when translocated into recipient cells, calculated as the percentage of transconjugants selected based on color or Em-resistance that were found to be edited. The mean ± SD is represented. **d)** Frequency of edition of MobA-BE per transconjugant, calculated as the percentage of Em-resistant transconjugants. The mean ± SD is represented. *, P < 0.05. **e)** Chromatograms obtained from the sequencing of transconjugants (T.) in which lacZ is targeted by TrwC-BE (left) or ermB is targeted by MobA-BE (right). The wild type (WT) sequence is provided as a reference. Edited nucleotides are marked with asterisks. The effect of the mutation in the protein product is indicated at the top: Syn. generates a synonymous codon, Stop introduces a stop codon, Start regenerates the mutated start codon.

Similar to the results observed upon electroporation of BE, many of the cells analysed contained a mixture of edited and WT sequences, and we noticed again a preference for editing the C that does not generate a stop codon in *lacZ* (**Fig. 4d**). For *ermB*, we identified 4% of double editing events, simultaneously at the desired position 12 after the PAM, and at position 16, close to the canonical editing window (**Fig. 4e**).

In conclusion, our findings demonstrate that relaxase-BE fusions can also be translocated via conjugation into a recipient cell, where they can refold and regain their gRNA-guided editing activity.

## Discussion

Targeted genetic modification of bacteria is fostering an increasing number of biotechnological and biomedical applications. The CRISPR-Cas technology has boosted the field, making targeted mutations a routine task for many model microorganisms (36). This technology has allowed metabolic engineering of different bacteria such as *E. coli* (37)*, Clostridium spp.* (38) or *Cyanobacteria spp.* (39), improving their use as cell factories. The system has also been used for biomedical research to study different pathogens such as *Mycobacterium tuberculosis, Yersinia pestis* o *Klebsiella pneumoniae* (36). CRISPR-Cas can also target specific bacterial or plasmid populations, enabling their use as antimicrobials (12, 40, 41). In spite of its success, the technology still faces significant limitations. A critical step to accomplish a genetic modification is the delivery of the endonuclease, gRNA, and template DNA to the target cell. Electroporation of plasmid DNA can be used to transform a wide range of bacteria, but protocols have to be fine-tuned for each new target strain, success is not guaranteed, and efficiencies are often very low. Phages have also been used for CRISPR-Cas delivery into bacteria, but the narrow specificity or bacterial resistance to infection pose significant limitations (42). Bacterial conjugation, a natural means of delivering DNA complexes into a wide range of bacteria, has been used to introduce the CRISPR-Cas genetic system into different species (23, 43, 44), avoiding the need for electroporation. Still, ensuring expression of the endonuclease in the recipient cell can be challenging in poorly characterized microorganisms, while overexpression can lead to toxicity and off-target activity (45).

In this work, we take advantage of the fact that the conjugative machinery delivers a nucleoprotein complex to the recipient cell. During conjugation, a relaxase protein is recognized by the T4SS and translocated covalently attached to the transferred DNA molecule. The rationale was to fuse the Cas endonuclease to the conjugative relaxase, so that the Cas protein itself is delivered *in vivo* to any bacteria that can be reached by the T4SS, bypassing the need for transcription and translation of a *cas* gene. Since the relaxase carries along the covalently attached DNA strand of our choice, with the only requisite of encoding a short *oriT*, the transferred DNA could encode the gRNA as well as template DNA for recombination-mediated seamless genetic modifications.

To show proof of concept of this strategy, we have selected the Cas endonuclease Cas12a. Cas12a has emerged as an interesting alternative to Cas9. Its different endonuclease architecture and mode of action makes it ideal for multiple events of genome editing. Its smaller size could also be crucial for translocation through the T4SS channel. For the relaxase moiety we chose TrwC, the well-characterized relaxase from the broad-host-range plasmid R388 (19), for several reasons. TrwC has been successfully fused to other polypeptides without losing its relaxase activity (46, 47). It can be translocated into the recipient cell either alone or linked to the DNA, and its activity in the recipient cell has been directly demonstrated (48, 49). Furthermore, TrwC can be translocated, by its own or by heterologous T4SS (31, 46, 48), into a wide range of organisms, including most proteobacterial species (50), Gram-positive bacteria (51), cyanobacteria (52), and even mammalian cells (53).

The TrwC-Cas12a protein, once constructed was evaluated for both stability and functionality. Although the stability of the protein was low, we were able to detect the full-size fusion protein. This protein showed RNA-guided endonuclease activity in bacteria, where it killed 99.9% of cells containing the target (**Fig. S1**). For TrwC to function in the recipient cell, it must partially unfold during secretion, and in fact, the inclusion of an unfolding resistant domain was shown to preclude protein translocation (54). The detection of Cas12a activity in the recipient cell after translocation of TrwC-Cas12a (**Figs. 1 and 2**) confirms that the complete fusion protein, despite its size of 263 kDa, can be translocated through the T4SS. We therefore conclude that there are no unfolding-resistant domains in Cas12a interfering with the translocation of the fusion protein. the detection of Cas12a activity in the recipient cell after translocation also demonstrates that the CRISPR endonuclease recovers its native structure after unfolding.

We have demonstrated the application of TrwC-Cas12a, delivered via conjugation, for targeted mutagenesis. This was achieved by translocating the fusion protein into a recipient strain that encodes *sacB*, and subsequently selecting *sacB* mutants. Sucrose-resistant transconjugants were only obtained when the appropriate gRNA was also expressed in the recipient. Analysis of the DNA sequence of the *sacB* region revealed small deletions at the expected TrwC-Cas12a cleavage site (**Fig. 1d**). This provides unequivocal evidence that the mutations were triggered by the RNA-guided endonuclease activity of TrwC-Cas12a.

The analysis of the *sacB* mutants has enabled us to observe the mutation pattern resulting from Cas12a-induced DSB repair in bacteria, which is not well documented in the literature. Absence of amplification of *sacB* in some mutants suggests the loss of the *sacB* locus by large deletions, as previously described after Cas9 cleavage in the chromosome of *E. coli* (25) as well as after Cas12a cleavage (55). The most common mutation consists of small 1-4 nt deletions at the cleavage site (**Fig. 1d**, TC 1-6). We have demonstrated that these deletions are solely the result of DNA cleavage by Cas12a (**Fig. 1d**, TF1-3), ruling out a possible effect of the relaxase moiety. To our knowledge, such indels have not been previously reported after cleavage by any CRISPR-Cas system in the chromosome of *E. coli*. However, a similar pattern of Cas12a-induced mutations has been identified in other prokaryotes, such as *Amycolatopsis mediterranei* (56). The introduction of small deletions proves to be a useful strategy for gene knockout.

To develop a delivery system capable of modifying wild-type recipient bacteria, it is necessary to introduce not only the Cas endonuclease, but also the gRNA. We used the conjugatively transferred DNA strand to encode the gRNA, so that both the nuclease and the gRNA would be provided from the donor cell. In this case, to our surprise, we observed an increase in the efficiency of Cas12a, as inferred by the observed lethality in the transconjugants, only when both the nuclease and the gRNA were translocated, resulting in the introduction of a DSB (**Fig. 2b**). This effect was not observed in the previous assay when the gRNA was encoded in the recipient cell. We hypothesize that the increase in Cas12a cleavage efficiency may be attributed to the physical proximity of the nuclease and the gRNA-encoding DNA in the translocated nucleoprotein complex.

We have explored the potential of using conjugatively delivered TrwC-Cas12a::DNA complexes to introduce the template DNA, carrying the desired mutation, into the target bacteria, in order to accomplish seamless targeted mutations by homologous recombination. We constructed a mobilizable suicide plasmid that carries a homologous recombination template of the *sacB* gene carrying a point mutation in the PAM sequence, which generated a premature STOP codon. We then mobilized this plasmid, either by by TrwC or TrwC-Cas12a into a recipient cell carrying the targeting gRNA or without it. Our goal was to obtain colonies that were sucrose-resistant and Cm-sensitive, which is the expected phenotype for the double-recombinants incorporating the mutation (SI Appendix **Fig. S2**). Upon analyzing the colonies from the different matings, we found that even though the number of sucrose-resistant colonies obtained was similar under the different conditions tested, the ratio of edited cell was significantly higher when both TrwC-Cas12a and the gRNA were present. Over 40% of the colonies had incorporated the desired mutations, while when TrwC was used or in the absence of the gRNA, this fraction dropped to one in 15 (**Fig. 2c**). In a previous work, it was shown that fusing Cas9 with the relaxase VirD2, increased the rate of homologous recombination up to 5.5-fold in plant cells, compared to the efficiency of Cas9 alone. In this experiment, the fusion protein, covalently attached to a homologous repair template, was bombarded into rice calli. The study suggests that the proximity of the repair template to the DBS, due to its covalent attachment to the relaxase, enhances the efficiency of homologous recombination. Therefore, it is plausible that our fusion protein may exhibit a similar effect (57).

Our results demonstrate a proof of concept for the use of bacterial conjugation to deliver in vivo relaxase-Cas fusion proteins, along with the covalently bound DNA template, into a recipient cell. However, the editing efficiency is currently too low for practical application, with only about 1 in 50,000 transconjugants resulting in edited, sucrose-resistant colonies. Thus, this approach would require selection of edited cells, which is rarely possible outside model systems like the one used in this study.

CRISPR-Cas editing strategies in bacteria typically hinge on the principle that unless the desired mutation is incorporated, the Cas nuclease’s cleavage of the target sequence will result in the death of the target bacteria. This allows for the introduction of mutations without the need for a selectable phenotype. The cleavage of a target sequence in the *E. coli* chromosome will only result in bacterial death if the cleavage is rapid enough to prevent homology-directed repair with an intact copy of the chromosome (25). While the delivery of the TrwC-Cas12 protein through the T4SS was not sufficient to kill target bacteria, likely because the T4SS most likely delivers a small number of Cas12 proteins to a single recipient, the TrwC-Cas12a:DNA complex encoding the gRNA did induce about 90% lethality in the transconjugants. This opens up the possibility of obtaining mutations without prior selection. A Cas protein must find two molecular partners in the recipient cell: the gRNA and then the target DNA. This lengthy search affects the speed of cleavage (58). By linking covalently Cas to the DNA encoding the gRNA, the first part of this search is likely shorter significantly. Future strategies that leverage this fact and increase the amount of Cas proteins delivered, may lead to a more efficient cleavage which provides the typical counter-selection mechanism for CRISPR-Cas genome editing strategies in bacteria.

Counter-selection of the wild-type genotype is only necessary when the editing efficiency is low, as is often the case with commonly used recombineering strategies. Novel strategies have been developed that do not depend on the introduction of double strand breaks to perform targeted genetic modifications. These strategies, such as base editing or prime editing, appear to work at high efficiencies (5). To demonstrate the versatility of our delivery system, we have also tested the ability to send base editing tools via bacterial conjugation. We created a fusion protein that consists of a conjugative relaxase and BE, which is itself a triple fusion between the cytidine deaminase, the dead version of Cas12a, and the UGI. This triple fusion (referred to here to as BE) has been previously tested in eukaryotes (28), but not in bacteria. To the N-terminus of this fusion protein, we added either the conjugative relaxase TrwC or the MobA relaxase of RSF1010. The latter is another well-characterized relaxase known to be active in recipients after translocation through different T4SS (59, 60). In addition, MobA uses the T4SS of plasmid RP4, which is known for its promiscuity, especially among distantly related donor/recipient pairs (61–63), thus increasing the range of potential recipient cells.

We have designed two systems to detect base editing, based on either a loss (*lacZ*) or gain (*ermB*) of function. These systems were initially used to confirm the BE activity of the fusion proteins in bacteria by electroporation (**Fig. 3**). All fusion proteins were found to be active at levels comparable to BE alone. We then confirmed their activity in recipients upon conjugation **(Fig. 4**). By using both selection systems, we verified that the vast majority of the selected transconjugants in these experiments had incorporated the expected mutation. BE as a fusion behaves similarly to its standalone form, with the same limitations, such as lower efficiency when editing C that are found after a G (as is the case of the C that generates the stop codon in *lacZ*), or occasional editing outside of what is strictly defined as the editing window (64). However, the frequency of edition in the absence of selection is around 0.1% of transconjugants. Even though this frequency is 50-fold higher than DSB-induced homology repair, it is still too low to be useful for the generation of mutations without selection. Future work will address the improvement of the efficiency for this purpose.

Finally, it is worth noting that these large multidomain proteins, which are bigger than 250 kDa, can be transported through the T4SS. They can then refold and regain activity in recipient cells, as evidenced by the base editing activity detected in these cells. These findings suggest that unfolding through the T4SS is not extensive. Indeed, recent studies have revealed that when TrwC is covalently attached to a ssDNA and translocated through the T4SS, the complex undergoes seven discrete steps of co-translocational unfolding, each stage representing a different translocation intermediate at various stages of unfolding (65). Therefore, the T4SS could serve as an effective delivery channel for proteins that would not be able to regain activity if they were completely unfolded during translocation, as is the case with T3SS-mediated translocation (66).

In summary, we present a proof of concept demonstrating that bacterial conjugation can be utilized to deliver active Cas endonucleases and base editors to recipient bacteria. This method eliminates the need for endonuclease expression in the recipient cell. The nucleoprotein complex can include any DNA of interest, which could encode the gRNA plus template DNA for homologous recombination. The delivery system is flexible enough to accommodate different relaxases and genetic editing tools, as evidenced by the successful delivery of a quadruple fusion base editor system. Considering the ability of broad-host-range conjugative systems to reach almost all Gram-negative, and even Gram-positive bacteria, conjugation could be widely used for the targeted genetic modification of prokaryotes, especially wild-type strains which are difficult to transform, and poorly characterized genera for which gene expression tools are underdeveloped (51). Even more, we have reported that T4SS from human pathogens can also translocate conjugative relaxases complexed with DNA to human cells (31, 67). Recently, the use of extracellular contractile injection systems (similar to Type VI secretion systems) has been described for this very purpose, demonstrating their ability to deliver proteins such as Cas9 and base editors into mammalian target cells (68). Current experiments are exploring the potential for Relaxase-Cas proteins to be delivered from bacteria and guide site-specific mutagenesis of human genes.

## Materials and Methods

### Bacterial strains and plasmids

Bacterial strains used in this work are listed in **SI AppendixTable S2.** *Escherichia coli* strains were grown at 37°C in Luria-Bertani (LB) broth, supplemented with agar for solid culture. Details of the strains used for the different experiments and the construction of the *E. coli* screening strain FD3 are explained in **SI Appendix Materials and Methods**.

Bacterial plasmids used in this work are listed in **SI Appendix Table S3**. Plasmid constructions and primers are detailed in **SI Appendix Materials and Methods and in SI Appendix Table S4**. **Table 1** summarizes the main features of key plasmids used in this work.

### Mating assays

Mating assays were performed as described in (69) with the modifications detailed in **SI Appendix Materials and Methods**. Briefly, cultures of donor and recipient strains, induced as indicated to express the Relaxase-Cas fusion proteins and/or the gRNA, were mixed on LB plates and mating plates were incubated for 1-3 h, depending on the experiment, at 37°C. Dilutions were plated on selective media for donors, recipients, transconjugants, or edited colonies, as required. Plates were supplemented with the specific inductor when indicated. Conjugation frequencies are expressed as the number of transconjugants per donor cell.

### Measurement of Cas12a cleavage activity in bacteria

In order to detect Cas12a cleavage activity in prokaryotic cells, we measured SOS induction and DSB-induced lethality.

#### SOS response assay

In order to detect induction of the SOS response upon translocation of TrwC-Cas12a, the plasmid pSOS was introduced by electroporation into the recipient strains MG1655 or MG1655*::sacB*. After the matings, Green Fluorescent Protein (GFP) levels were detected directly on the conjugation plates in an Azure Biosystems c400 imaging system. Next, conjugation was stopped by introducing the filter in 2 ml LB broth. 100 µl were added on a 96 well black flat microtiter plate. GFP signal (excitation filter: 475 nm and emission filter: 515 nm) and bacterial cell density (OD 600nm) were measured with a TECAN infinite M200 Pro plate reader.

#### Lethality assay

100 ng of each plasmid encoding Cas12a and gRNA (*lacZ* or *sacB*) were electroporated into D1210, and cells were plated on antibiotic containing media supplemented with IPTG 500 µM for gRNA expression. aTc 200 ng/ml for induction of *cas12a* expression was added when indicated. Lethality was measured by comparing the number of CFU with/without induction or with/without target for the gRNA. In order to measure Cas12a-induced lethality upon translocation into the recipient cell, we compared the number of transconjugants obtained under these conditions.

#### RNA-guided mutations in sacB

Plasmids encoding the nuclease and *sacB* gRNA were electroporated into MG1655*::sacB*, and cells were plated on antibiotic containing media supplemented with IPTG 500 µM and 1% sucrose to counterselect *sacB* activity. To detect mutations upon translocation of TrwC-Cas12a into the recipient MG1655:*sacB* strain (strain FD3, **SI appendix Table S2**) the matings were directly plated on 1% sucrose-containing plates. Sucrose-resistant transconjugants were directly picked for PCR amplification of the *sacB* region using sacB_F and sacB_R primers (**SI Appendix Table S4**). The size of the amplicon was checked by agarose gel electrophoresis. Amplicons were purified and their DNA sequence was determined (STAB VIDA). Sequence alignments were performed with BioEdit Sequence Alignment Editor.

### Measurement of seamless editing rate

After matings, sucrose-resistant colonies were replicated in Cm-containing plates to discard integrants (Cm-resistant). Then, a fragment of 357 bp from *sacB* containing the PAM region was amplified from the Cm-sensitive transconjugants using the oligonucleotides sacB_F and sacB_HR_R (**SI Appendix Table S4**). Amplicons were purified and their DNA sequence was determined (STAB VIDA). Sequence alignments were performed with BioEdit Sequence Alignment Editor.

The editing rate was calculated as the number of Cm-sensitive colonies incorporating the desired mutation (stop codon) divided by the number of sucrose-resistant colonies analyzed in each assay.

### Analysis of BE-edited colonies

To test the activity of BE and its fusions by electroporation, 100 ng of plasmids containing BE, relaxases or relaxase-BE fusions and 100 ng of pgRNA-lacZ_BE_ were coelectroporated in D1210. Cells were plated on antibiotic containing media supplemented with X-Gal 60 μg/ml, IPTG 500 µM and aTc 200 ng/ml (except in the case of pTrwC, where aTc is not added). Light-colored transformants (visually detected) were passaged 1 to 3 times to fix the edition. Afterwards, the *lacZ* region was PCR-amplified using lacZ_F and lacZ_R primers (**SI Appendix Table S4**). For assays targeting plasmid-borne *ermB*\*, 100 ng of the same plasmids were electroporated in DH5α containing plasmids pErmB* and pgRNA-ErmB_BE_. Transformants were selected on Em Ap Ara plates and grown for 48 h. Em-resistant colonies were replicated again in the same medium and picked for PCR amplification of the *ermB* region with araBAD_F and pBAD33_seq_R primers (**SI Appendix Table S4**). The size of the amplicons was checked by agarose gel electrophoresis. All PCR products were purified, and DNA sequence was determined (STAB VIDA). When the ratio between Em-resistant and total transformants was needed, transformants were selected in media containing only antibiotics for plasmid selection, and then transferred to Em Ara Ap plates. Then, the percentage of Em-resistant colonies was calculated.

To verify edition following the translocation of relaxases and their fusions by conjugation, transconjugants were plated on selective media. In the case of *lacZ*, the medium contained the antibiotics needed for transconjugant selection, along with X-Gal and IPTG. To detect edition of *ermB*, transconjugants were selected on two types of media: one with the necessary antibiotics for their selection, to obtain the total number of transconjugants, and another with Em Ap Ara to selectively isolate and count those resistant to Em. Selection of the colonies, passages, PCR amplification of the target genes and sequencing was done in the same way as in the case of electroporation.

### Western Blot

Total protein extracts were obtained as described in (70). *E. coli* D1210 cells containing plasmids pTrwC, pCas12a, pTrwC-Cas12a, pBE or pTrwC-BE were grown overnight. The cultures were diluted 1:20 and induced with IPTG 500 µM or aTc 200 ng/ml for 3 h. 1 ml of each culture was collected, centrifuged, and resuspended in 1/10 volume of 2xSDS-gel loading buffer. Samples were stored at -20°C for at least overnight. Samples were boiled for 5 min and loaded on 9% acrylamide SDS-PAGE gels. After the run, the gels were transferred to nitrocellulose membranes. Primary antibody (anti-TrwC (71)) and secondary antibody (Anti-Rabbit IgG IRDye®800CW, Li-Cor) were used at 1:10,000 dilution. Detection was performed with an Odyssey Clx. NZYColour Protein Marker II and PageRule Plus Prestained Protein Ladder (Thermo Fisher) were used as molecular weight marker.

### Statistical analysis

A Student’s t-test was employed to identify statistically significant discrepancies between the means of three or more independent results. When comparing more than two groups, a one-way ANOVA was utilized. The non-parametric Mann-Whitney U test was used for data that did not follow a Gaussian distribution. All statistical analyses were conducted using GraphPad Prism 8 software.

## Acknowledgments

Work in ML lab is supported by grants PID2020-117956RB-I00 and PDC2021-120967-I00_MCIN/AEI/10.13039/501100011033_UE Next GenerationEU/PRTR from the Spanish National Research Agency (Ministry of Science and Innovation). DB was supported by the European Research Council [677823]; European Research Council [101044479]; Agence Nationale de la Recherche [ANR-10-LABX-62-IBEID]. AF-G was a recipient of a predoctoral appointment from the University of Cantabria.

## Author Contributions

DLG-H, AF-G and FD performed the experiments. ML and DB designed the work. ML, DB, DLG-H and AF-G analysed and interpreted the results, and wrote the manuscript.

## Classification

Biological Sciences

Microbiology.

