## Supplementary information for "In vivo delivery of functional Cas:DNA nucleoprotein complexes into recipient bacteria through a Type IV Secretion System"

### Material and Methods

#### *Bacterial strains and culture media*

Bacterial strains are listed in **Table S2**. *E. coli* strains DH5α (T1 phage resistant) and D1210 were used for routine cloning and as donor and recipient strains in matings. D1210 was used also for the lethality assay in prokaryotic cells. MG1655 and FD3 were used as recipient cells in SOS response and *sacB* mutants detection assays. DH5α*pir* and β2150 were used as donor and recipient strains respectively for mobilization of the homologous recombination cassette. *E. coli* 4644 was used as host for recombinant plasmids based on pOSIP-KL and was grown at 30°C. Selective media included the following antibiotics at the indicated concentrations: chloramphenicol (Cm) 20 µg/ml; ampicillin (Ap) 100 µg/ml; kanamycin monosulfate (Km) 20 µg/ml; streptomycin (Sm) 300 µg/ml; gentamicin sulfate (Gm) 10 µg/ml; and nalidixic acid (Nx) 20µg/ml. aTc (anhydrotetracycline hydrochloride), IPTG (Isopropyl β- d-1-thiogalactopyranoside), X-Gal (5-bromo-4-chloro-3-indolyl-β-D-galactopyranoside) and diaminopimelic acid (DAP) were supplied at a concentration of 200 ng/ml, 500 µM, 40 µg/ml and 0.3 mM respectively. Plates containing 1% sucrose were used for the selection of *sacB* mutants.

#### *Construction of the* E. coli *screening strain FD3*

The strain FD3 was constructed through integration of plasmid pFD148 (**Table S3**) at the phage lambda *attB* site of strain MG1655. Plasmid pFD148 is an integrative vector based on pOSIP-KL, which carries the *yhhX* target sequence on the non-template strand between a constitutive promoter and the *mCherry* reporter gene followed by a *sacB* counter-selection marker. Plasmid pFD148, containing the lambda integrase, lambda *attP* site and a kanamycin resistance cassette, was integrated at the lambda *attB* site in the chromosome of *E. coli* MG1655. The integration was verified by PCR and the backbone was flipped out using the pE-FLP plasmid able to recombine FRT sites flanking the backbone as previously described (1). pE-FLP was then cured through serial restreaks on LB plates, leading to the strain FD3. For the sake of clarity, we will refer to this strain as MG1655::*sacB*.

#### *Plasmid constructions.*

PCR amplifications for cloning procedures were performed with Phusion High-Fidelity DNA Polymerase (Thermo Scientific™) or with Kapa HiFi DNA polymerase (Kapa biosystems). Plasmids used in this work and the oligonucleotides used for each construction are listed in **Table S3** and **S4** respectively**.** Those plasmids mentioned in the text have been renamed for clarity (Table S3, Name in the manuscript column). Analytical PCR reactions were performed with Kapa Taq polimease (Kapa biosystems). Plasmids pAF18 (pMobA-BE), pAF21 (pErmB*), pAF21 (pMobA), pFD148, pLG14 (pCas12a), pLG22 (pTrwC), pLG24 (pTrwC-Cas12a), pLG27 (pHR_oriT), pLG29 (pgRNA-sacB-oriT), pLG32 (pBE), pLG33 (pTrwC-BE), were constructed by the isothermal assembly method (2). pLG15 (pgRNA-lacZ) and pLG19 (pgRNA-sacB) were constructed by standard restriction cloning (3).

- pFD148: A 1873-pb fragment containing *sacB* was amplified from total DNA of *Bacillus subtilis* 168 and introduced upstream *mCherry* reporter gene by PCR amplification of plasmid pFD145 followed by isothermal assembly.
- pCas12a was constructed assembling *cas12a*-NLS-3Ha amplified from pY010 and PCR-amplified pBBR6, leaving *cas12a*-NLS-3Ha downstream the lactose promoter.
- pTrwC was constructed assembling *trwC* amplified from pHP159 and PCR-amplified pBBR6. *trwC* was inserted downstream the lactose promoter.
- pTrwC-Cas12a was constructed assembling 4 amplicons: P*tet* promoter was amplified from the pOSIP-CO-RBS-library-dCas9 plasmid. *trwC* sequence without stop codon was amplified from pAA12. pCas12a was amplified in 2 fragments, without amplifying the lactose promoter. *trwC* was assembled upstream *cas12a* in frame with its start codon and the P*tet* fragment was inserted upstream the fusion gene.
- pHR_oriT was constructed assembling a *sacB* homologous recombination cassette under the control of a P*tac* promoter with PCR-amplified pSW27. The 5' 430 bp of sacB were amplified from strain FD3 in two PCR reactions, in order to introduce a mutation in the PAM sequence leading to an early stop codon. The P*tac* promoter was added to the oligonucleotides upstream the homologous recombination cassette.
- pgRNA-lacZ and pgRNA-sacB were constructed by inserting into pUC8 and pUC18 respectively an *Eco*RI-*Hin*dIII fragment containing gRNAs for *lacZ* and *sacB* respectively (synthesized as gBlocks; ATG:Biosynthetics). The DNA sequences are shown in Table S4.
- pgRNA-sacB-oriT was constructed by adding an R388 *oriT* sequence into the pgRNA-sacB plasmid by isothermal assembly.
- pTrwC-BE was constructed assembling *dCpf1-BE* amplified from pCMV-dCpf1-BE and PCR-amplified pTrwC-Cas12a with *trwC* sequence but without *cas12a* sequence, which was replaced by *dCpf1-BE*, placed after *trwC* without stop codon.
- pBE was constructed assembling *dCpf1-BE* amplified from pCMV-dCpf1-BE and PCR-amplified pTrwC-Cas12a without *trwC-cas12a*, which was replaced by *dCpf1-BE*.
- pMobA-BE was constructed assembling *mobA* from RSF1010, without stop codon, amplified from pLG04, and PCR-amplified pTrwC-BE without *trwC* sequence, which was replaced by *mobA*.
- pMobA was constructed assembling *mobA* from RSF1010 amplified from pLG04 and PCR-amplified pTrwC-BE without *trwC-dCpf1-BE* sequence, which was then substituted with *mobA*.

- pErmB* was constructed assembling *ermB*, amplified from pAM1 and modified to change its ATG start codon to ACG, and the PCR amplified pBAD33 vector, so that *ermB* was inserted downstream the P*bad* promoter. A Shine Dalgarno sequence and a PAM sequence were also introduced in the 5' region of the gene.
- pgRNA-lacZ$_{BE}$ and pgRNA-ErmB$_{BE}$ were constructed by inserting into pUC18 EcoRI-HindIII fragments containing gRNAs for *lacZ* (pgRNA-lacZ$_{BE}$) or *ermB* (pgRNA-ErmB$_{BE}$ ). The DNA sequences are shown in Table S4.

**Mating assays**

In matings to test for TrwC-Cas12a and TrwC-BE function in conjugation, D1210 and DH5α were used as donor and recipient bacteria respectively. For MobA-BE, the donor was S17.1. Induction conditions were as follows: overnight cultures of donor and recipient strains were diluted 1/20 and grown for 3 h in the presence of IPTG 0.5 mM for TrwC or aTc 200 ng/ml for the rest. 100µl were mixed with 100 µl of overnight recipient culture and placed for 1-3 h on cellulose acetate filter on a LB agar plate supplemented with the specific inducer.

For matings to test the mobilization of the pHR_oriT, *pir*+ strains DH5αpir and β2150 were used as donor and recipient respectively. The donor strain included plasmids pHR_oriT, R388*trwC*-, and either pTrwC or pTrwC-Cas12a.

For mating assays to test for Cas12a activity in recipient cells, D1210 was used as donor and MG1655 or MG1655::*sacB* as recipient strains. For mating assays to test for incorporation of site-specific mutation using a homologous recombination cassette, DH5α*pir* was used as donor and MG1655::*sacB* as recipient strain. In mating assays to detect transconjugants edited by BE, both the donor and the recipient were D1210 in the case of TrwC-BE, while for MobA-BE, the donor was S17.1 and the recipient was DH5α with the target plasmid. Overnight cultures of donor and recipient strains were diluted 1/20 and grown for 3 h in the presence of aTc 200 ng/ml (donor) and IPTG 500 µM (recipient). The mixture was placed on a LB agar plate supplemented with aTc 200 ng/ml and IPTG 500 µM. Transconjugants, integrants or recipients (depending on the assay) were also selected on plates supplemented with antibiotics. When indicated, 1% sucrose was added in order to select sucrose-resistant mutants, Em 200 µg/mL and arabinose 0.1% to select Em-resistant transconjugants, and X-Gal 60 µg/mL to screen for white colonies.

**Table S1. Control complementation assays with fusion proteins**

| Donor → recipient | Mobilizable plasmid | Complementing plasmid | Conjugation frequency[1] |
|---|---|---|---|
| **Complementation assays with relaxase-Cas fusion proteins[2]** | | | |
| **D1210→ DH5α $T1^R$** | R388*trwC-* | pTrwC | 5.48 x $10^{-1}$ ± 2.7 x $10^{-1}$ |
| | | pTrwC-Cas12a | 4.1 x $10^{-1}$ ± 2.6 x $10^{-1}$ |
| | | none | <1.8 x $10^{-8}$ |
| **pHR_oriT mobilization[3]** | | | |
| **DH5α*pir*→ β2150** | pHR_oriT R388*trwC-* | pTrwC | 6.5 x $10^{-1}$ ± 1.2 x $10^{-1}$ |
| | | pTrwC-Cas12a | 5.3 x $10^{-1}$ ± 1 x $10^{-1}$ |
| | | none | < 3.7 x $10^{-8}$ |
| **Complementation assays with relaxase-BE fusion proteins[3]** | | | |
| **D1210→ DH5α $T1^R$** | R388_TrwC- | pTrwC | 7.52 x $10^{-2}$ ± 3.86 x $10^{-2}$ |
| | | pTrwC-BE | 3.72 x $10^{-2}$ ± 1.21 x $10^{-1}$ |
| | RSF1010*mobA-* | pMobA | 8.35 x $10^{-1}$ ± 3.95 x $10^{-1}$ |
| | | pMobA-BE | 17.06 x $10^{0}$ ± 8.26 x $10^{0}$ |

[1] Transconjugants per donor
[2] Data represent the mean +/- SD of 4 independent assays
[3] Data represent the mean +/- SD of 3 independent assays

**Table S2. Bacterial strains**

| Strain | Relevant genotype | Reference |
|---|---|---|
| ***Bacillus subtilis 168*** | Contains *sacB* | (4) |
| ***Escherichia coli* 4644** | TransforMax$^{TM}$ Ec100D$^{TM}$ pir+ | Lucigen |
| ***E. coli* D1210** | *recA hspR hsdM rpsI lacI$^q$* | (5) |
| ***E. coli* DH5α *pir*** | *endA1 hsdR17 glnV44 thi-1 recA1 gyrA96 relA1 φ80dlacΔ(lacZ)M15 Δ(lacZYA-argF)U169 zdg-232::Tn10 uidA::pir+* | (6) |
| ***E. coli* DH5αT1 phage resistant** | *F- ϕ80lacZΔM15 Δ((lacZYA-argF)U169 recA1 endA1 hsdR17(rk−, mk+) phoA supE44 λ-thi-1 gyrA96 relA1 tonA* | (7) |
| ***E. coli* FD3** | MG1655::*mcherry::sacB* | This work |
| ***E. coli* MG1655** | F- lambda- *ilvG- rfb-50 rph*-1 | (8) |
| ***E. coli* S17.1** | SmR ; *F-* RP4-2-Tc::Mu *aph*::Tn7 recA | (9) |
| ***E. coli* β2150** | *ΔdapA::(erm-pir) thrB1004, pro, thi, strA, hsdS, lacZ ΔM15,(FΔ lacZ ΔM15 lacIq, traD36, proA+,proB+)* | (10) |

**Table S3. Plasmids used in this work**

| Plasmid | Name in the manuscript | Description | Reference |
|---|---|---|---|
| **pAA12** | | pHP159::*trwC-ralF TS*; Gm$^r$ | (11) |
| **pAA58** | | RSF1010K::*egfp*; Km$^r$ | (12) |
| **pAM1** | | *E. coli-Bifidobacterium* shuttle cloning vector; Ap$^r$ Em$^r$ | (13) |
| **pAF18** | pMobA-BE | pBBR6 *Δ*P*lac*:: P*tet mobA-dCpf1-BE*; Gm$^r$ | This work |
| **pAF21** | pMobA | pBBR6 *Δ*P*lac*::P*tet mobA*; Gm$^r$ | This work |
| **pAF20** | pErmB* | pBAD33:: P*bad ermB*(Start codon mutation)*; Cm$^r$ | This work |
| **pAF20-ATG** | pErmB | pBAD33:: P*bad ermB*; Cm$^r$ | This work |
| **pAF23** | pgRNA-ErmB$_{BE}$ | pUC18:: P*lac ermB*$_{gRNA}$; Ap$^r$ | This work |
| **pBAD33** | | Cloning vector; Cm$^r$ | |
| **pBBR6** | | Cloning vector derived from pBBR1-MCS; Gm$^r$ | (14) |
| **pE-FLP** | | One-Step cloning and integration plasmid | (1) |
| **pFD145** | | Contains *mCherry*; | (15) |
| **pFD148** | | pOSIP:*yhhX:mcherry:sacB* | This work |
| **pHP159** | | pBBR6::*oriT trwABC::pCMV eGFP*; Gm$^r$ | (16) |
| **pLG04** | | pAA58::*hyg*; Km$^r$ Hyg$^r$ | (12) |
| **pLG14** | pCas12a | pBBR6::*cas12a*; Gm$^r$ | This work |
| **pLG15** | pgRNA-lacZ | pUC8:: P*lac lacZ*$_{gRNA}$; Ap$^r$ | This work |
| **pLG19** | pgRNA-sacB | pUC18:: P*lac sacB*$_{gRNA}$; Ap$^r$ | This work |
| **pLG22** | pTrwC | pBBR6::*trwC*; Gm$^r$ | This work |
| **pLG24** | pTrwC-Cas12a | pBBR6 *Δ* P*lac*::P*tet trwC-cas12a*; Gm$^r$ | This work |
| **pLG27** | pHR_oriT | pSW27::P*tac::sacB** homologous recombination cassette; Cm$^r$ | This work |
| **pLG29** | pgRNA-sacB-oriT | pLG19 *oriT* | This work |
| **pLG32** | pBE | pBBR6 *Δ* P*lac*::P*tet dCpf1-BE*; Gm$^r$ | This work |
| **pLG33** | pTrwc-BE | pBBR6 *Δ* P*lac*:: P*tet trwC-dCpf1-BE*; Gm$^r$ | This work |
| **pLG38** | pgRNA-lacZ$_{BE}$ | pUC18:: P*lac lacZ*$_{gRNA}$; Ap$^r$ | This work |
| **pMTX808** | RSF1010_MobA$^-$ | pAA58::*ap* in *mobA*; Ap$^r$ Km$^r$ | (12) |
| **pOSIP-KL** | | One-Step cloning and integration plasmid | (1) |
| **pOSIP-CO-RBS-library-dCas9** | | Contains P*tet* | (17) |
| **pSU1186** | poriT | pUC18::*oriT*$_{R388}$ | (18)‼ |
| **pSU1445** | R388*trwC-* | R388::*Tn5tac1* in *trwC*; Km$^r$ Tp$^r$ | (19) |
| **pSW27** | | pSW23*::oriT*$_{R388}$+*ori*VR6K; Cm$^r$ | (10) |
| **pUC8** | pgRNA-Empty | Cloning vector; Ap$^r$ | (20) |
| **pUC18** | | Cloning vector; Ap$^r$ | (21) |
| **pY010** | | pcDNA3.1:: *hAscas12a*; Ap$^r$ | (22) |
| **pZA31-sulA-GFP** | pSOS | pZA31::*pSOS gfp*; Cm$^r$ | (23) |

**Table S4. Oligonucleotides used in this work**

| Purpose | Oligonucleotide sequence (5' a 3')[1] |
|---|---|
| **Construction of pFD148** | |
| **F310-pFD145-sacB-F** | CAATTAACAGTTAACAAATAA**AGGCATGCCTCGAGATGCATG** |
| **F311-pFD145-sacB-R** | CTCCTTTTTTATGTACGCA**TTATTTGTACAGCTCATCCATG** |
| **F312- sacB-N-ter** | CTGTACAAATAA**TGCGTACATAAAAAAGGAGACATGAACGATG** |
| **F313-sacB-C-ter** | CTCGAGGCATGCCT**TTATTTGTTAACTGTTAATTGTCCTTGTTC** |
| **Construction of pCas12a** | |
| **pBBR6_1440_F** | **GCGTTAATATTTTGTTAAAATTCG** |
| **pBBR6_1074_R** | **AGCTGTTTCCTGTGTGAAA** |
| **Cpf1:BID_pBBR6_F** | TGAGCGGATAACAATTTCACACAGGAAACAGCT**ATGACACAGTTCGAGGG** |
| **pLG14_Cpf1_R** | TTTAACGCGAATTTTAACAAAATATTAACGC**GGCATAGTCGGGGACA** |
| **Construction of pgRNA-lacZ** | |
| ***lacZ*$_{gRNA}$ gBlock** | GAATTCGTCAAAAGACCTTTTTAATTTCTACTCTTGTAGATCCGACCGCACGCCGCATCCAGCGCTGTCAAAAGACCTTTTTAATTTCTACTCTTGTAGATAAGCTT |
| **Construction of pgRNA-sacB** | |
| ***sacB*$_{gRNA}$ gBlock** | GAATTCGTCAAAAGACCTTTTTAATTTCTACTCTTGTAGATGGACAGCTGGCCATTACAAAACGGTCAAAAGACCTTTTTAATTTCTACTCTTGTAGATAAGCTT |
| **Construction of pgRNA-sacB-oriT** | |
| **pLG28_F** | **GTGCACACAGCCCAGC** |
| **pLG28_R** | **GCTGAACGGGGGGTTC** |
| **pLG28_oriT_1** | TACCGGATAAGGCGCAGCGGTCGGGCTGAACGGGGGGTTC**CTCATTTTCTGCATCATTGT** |
| **pLG28_oriT-402** | TTCGGTGTAGGTCGTTCGCTCCAAGCTG**GGCTGTGTGCACCCGCCTCGTCCTCCAAAAG** |
| **Construction of pTrwC** | |
| **pBBR6_F** | **GCGTTAATATTTTGTTAAAATTCGC** |
| **pBBR6_R** | **AGCTGTTTCCTGTGTGAAATTGT** |
| **pBBR6_TrwC_F** | GGAATTGTGAGCGGATAACAATTTCACACAGGAAACAGCT**ATGCTCAGTCACATGGTATTGA** |
| **pBBR6_TrwC_R** | TAACAAAAATTTAACGCGAATTTTAACAAAATA**TTAACGCTTACCTTCCGGCCTCCAT** |
| **Construction of pTrwC-Cas12a** | |
| **tetR** | **TTAAGACCCACTTTCACATTTAAG** |
| **ptet** | **TTTTGCCTCCTAACTAGGTCAT** |
| **ptet_TrwC** | TATCCGGAGGCATATCAAATGACCTAGTTAGGAGGCAAAA**ATGCTCAGTCACATGGTATTGA** |
| **TrwC_Cpf1** | TCACCTGATACAGGTTGGTAAAGCCCTCGAACTGTGTCAT**TGACCTTCCGGCCTCCAT** |
| **Cpf1_F** | **ATGACACAGTTCGAGGGCT** |
| **pLG14R** | **GCGTAGCACCAGGCGT** |
| **plg14_F** | **ACCAATAGGCCGACTGCGAT** |
| **plg14_2_R** | CGGATTAGAAAAACAACTTAAATGTGAAAGTGGGTCTTAA**TTAGGTGGCGGTACTTGGGTCG** |
| **Construction of pHR_oriT** | |
| **pW27_R2.0:** | **AGTTCTAGAGCGGCCGCCA** |
| **pSW27_F2.0** | **AGTGGATCCCCCGGGCT** |
| **pSW27_sac_F** | **AGCTCCACCGCGGTGGCGGCCGCTCTAGAA**CTGTCAATAGAAGTTTCGCCGA |

| | |
|---|---|
| **pSW27_ptac-sacb_R** | **TGAGAATTCCTGCAGCCCGGGGGATCCACT**TTGACAATTAATCATCGGCTCGTATAATGTGCGTACATAAAAAAGGAGACAT |
| **sacB_F** | **TGGGACAGCTGGCCATTACAA** |
| **sacb_R** | **TTGTAATGGCCAGCTGTCCCATTAGTCCAGGCCTTTTGCA** |
| **Construction of pTrwC-BE** | |
| **pLG24_BE_F** | **GCGTTAATATTTTGTTAAAATTCGC** |
| **pLG24_pTet_R** | **TTTTGCCTCCTAACTAGGTC** |
| **BE_pTet_F** | ATCCGGAGGCATATCAAATGACCTAGTTAGGAGGCAAAA**CCCAAGAAGAAGAGGAAAGTC** |
| **BE_R** | AACAAAAAATTTAACGCGAATTTTAACAAAATATTAACGC**ATGGTGATGGTGATGATGAC** |
| **Construction of pgRNA-lacZ$_{BE}$** | |
| ***lacZ*$_{gRNA}$ gBlock** | GAATTCGTTTCAAAGATTAAATAATTTCTACTAAGTGTAGATGCTAAATACTGGCAGGCGTTTCGGTTTCAAAGATTAAATAATTTCTACTAAGTGTAGATAAGCTT |
| **Construction of pMobA-BE** | |
| **pLG33_F** | **ATGAGCTCAGAGACTGGCCC** |
| **pLG24_pTet_3.0** | **TTTTGCCTCCTAACTAGGTCATTTG** |
| **MobA_33_F** | TCCGGAGGCATATCAAATGACCTAGTTAGGAGGCAAAA**ATGGCGATTTATCACCTTACGG** |
| **MobA_solo_R** | TAAAAAATGAGCTGATTTAACAAAAATTTAACGCGAATCTA**CATGCTGAAATCTGGCCCG** |
| **Construction of pMobA** | |
| **pLG33_F** | **ATGAGCTCAGAGACTGGCCC** |
| **MobA-33_F** | TCCGGAGGCATATCAAATGACCTAGTTAGGAGGCAAAA**ATGGCGATTTATCACCTTACGG** |
| **MobA-33_R** | ATGTGGGGTCCACAGCCACTGGGCCAGTCTCTGAGCTCAT**CATGCTGAAATCTGGCCCG** |
| **pLG24_pTet_3.0** | **TTTTGCCTCCTAACTAGGTCATTTG** |
| **Construction of pErmB*** | |
| **pBAD33_MCS_F** | **GAGCTCGGTACCCGGGG** |
| **pBAD33_MCS_R** | **GAATTCGCTAGCCCAAAAAAAC** |
| **Erymut_F** | GTTTCTCCATACCCGTTTTTTTGGGCTAGCGAATTC*TTTAGGAGTAAATAAC***GAACGAGAAAAATATAAAACACAGTC** |
| **Erymut_R** | TGCAGGTCGACTCTAGAGGATCCCCGGGTACCGAGCTC**TTACTTATTAAATAATTTATAGCTATTGAAAAG** |
| **Construction of of pgRNA-sacB$_{BE}$** | |
| ***ermB*$_{gRNA}$ gBlock** | **GAATTCGTTTCAAAGATTAAATAATTTCTACTAAGTGTAGATGGAGTAAATAACGAACGAGAAAAGTTTCAAAGATTAAATAATTTCTACTAAGTGTAGATAAGCTT** |
| ***sacB* region amplification for transconjugants analysis** | |
| **SacB_F** | **CTACCGCACTGCTGGCAG** |
| **SacB_R** | **GATGCTGTCTTTGACAACAG** |
| ***sacB* region amplification for edited cells with the HR cassette** | |
| **SacB _F** | **CTACCGCACTGCTGGCAG** |
| **SacB_HR_R** | AGCTCCACCGCGGTGGCGGCCGCTCTAGAA**CTGTCAATAGAAGTTTCGCCGA-** |
| ***lacZ* region amplification for edited cells** | |
| **lacZ_F** | **CGGCGTTTCATCTGTGGTG** |
| **lacZ_R** | **CCGGCTGCGGTAGTTCAG** |
| ***ermB* region amplification for edited cells** | |
| **araBAD_F** | **GATTAGCGGATCCTACCTG** |

| | |
|---|---|
| **pBAD33_seq_R** | **CATGGGGTCAGGTGGG** |

[1] For plasmids constructed by isothermal assembly, nucleotides annealing to the template during PCR amplification are shown **in bold**. For plasmids constructed by classical recombinant DNA techniques, *Eco*RI and *Hin*dIII restriction sites used for cloning are underlined. See text for more details.

**Figure S1. Validation of TrwC-Cas12a fusion protein**

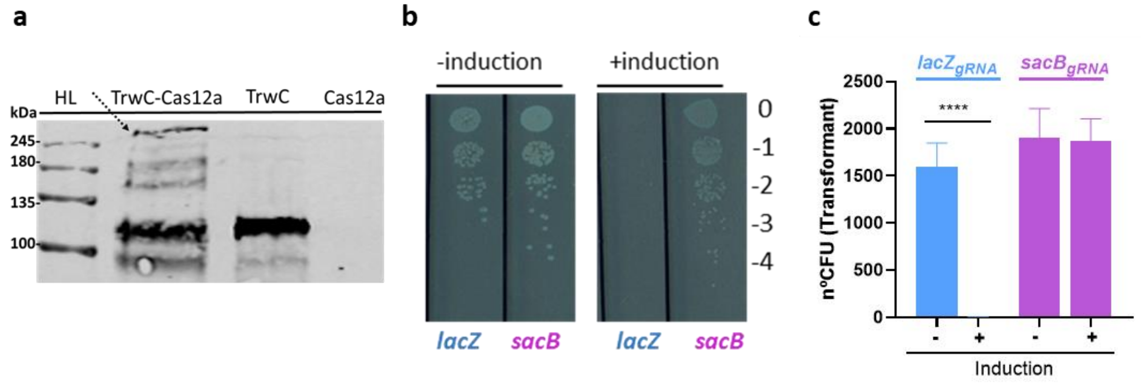


**a)** Stability of TrwC-Cas12a. A Western Blot was performed using an anti-TrwC antibody. D1210 cell lysate was used as negative control (-). TrwC, Cas12a and TrwC-Cas12a correspond with D1210 lysates carrying pTrwC, pCas12a and pTrwC-Cas12a respectively, after 3 hours of induction with IPTG (for pTrwC and pCas12a) or aTc (for pTrwC-Cas12a). The expected sizes for the proteins are: TrwC 108kDa, Cas12a 187 kDa, TrwC-Cas12a 263 kDa. Left lane, molecular weight marker. Black arrows indicate 245 kDa (top) and 100 kDa (bottom) bands. The dotted arrow indicates full-size TrwC-Cas12a protein. **b)** Nuclease activity of TrwC-Cas12a. D1210 coelectroporated with plasmids pTrwC-Cas12a and either pgRNA-lacZ or pgRNA-sacB were plated under non-induction or induction (aTc) conditions for the expression of *trwC-cas12a*. 10 µl of each dilution (0 to -4) were plated. **c)** Number of transformants after coelectroporation of *E. coli* D1210 with pTrwC-Cas12a and the plasmids encoding the indicated gRNA, under conditions of induction (+) or non-induction (-) of *trwC-cas12a* expression with aTc. Data correspond with the mean of at least 3 independent assays. ****, P<0.0001

**Figure S2.**

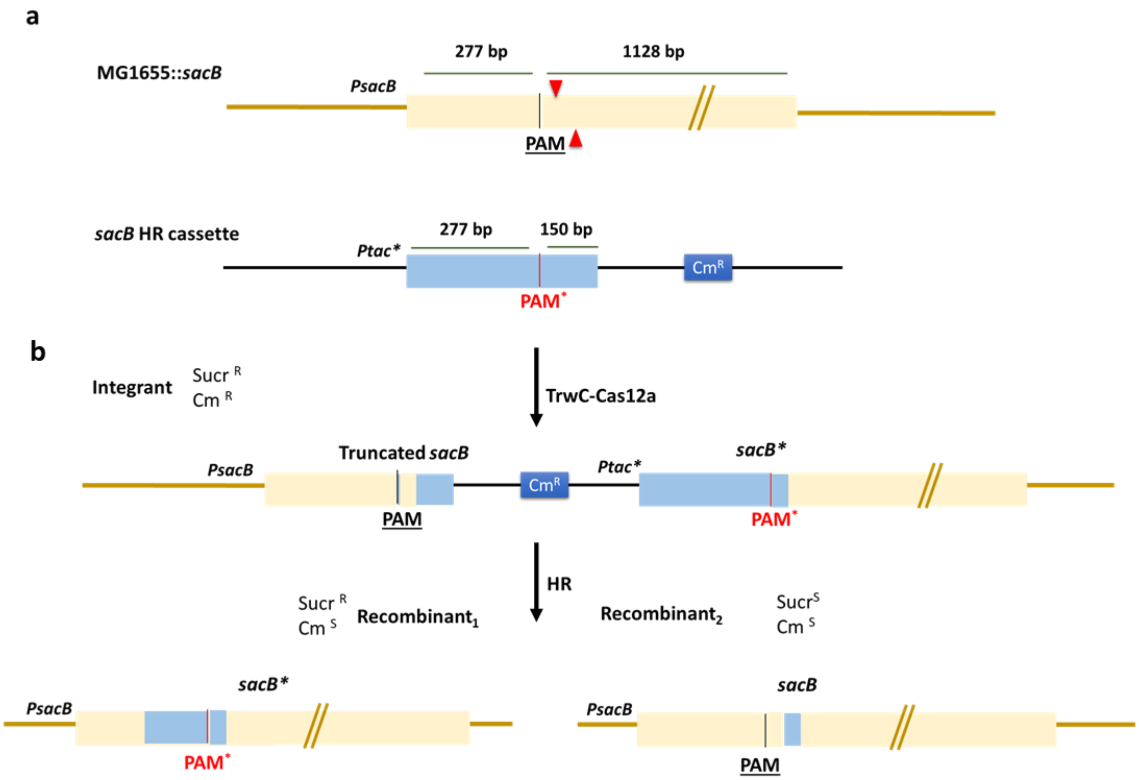


**Figure S2. Introduction of seamless mutations using a HR cassette. a)** Map of the *sacB* gene and the HR cassette. This cassette contains 430 bp of the 5' region of the *sacB* gene present in MG1655::*sacB* strain. This region includes the sequence targeted by the gRNA encoded by plasmid pgRNA-sacB. The PAM sequence in the homologous recombination cassette was mutated to generate an early STOP codon in the gene (TAA). Consequently, incorporation of the mutation also abolishes the PAM sequence, preventing other Cas12a-mediated cleavage events. The site of the mutation was surrounded by a left homology arm of 277 bp and a right homology arm of 150 bp**.** The red arrowheads mark the sites of Cas12a cleavage. **b)** TrwC-Cas12a mediated edition. Cas12a cleavage in the chromosomal *sacB* copy would generate a DSB which will be repaired by the HR pathways of the cell. The resulting integrants would carry a *sacB* copy with the wild type PAM and the right arm of the HR cassette (thus encoding a truncated *sacB*), and a second *sacB* copy with the mutated PAM (red vertical bar,) and the full *sacB* sequence. The second recombination could occur 5' from the PAM sequence, generating a recombinant (Recombinant1) with a *sacB* copy containing the mutated PAM (left), or it could occur 3' from the PAM sequence, generating the recombinant2, with a non-edited *sacB* copy.

**Figure S3. Stability of TrwC-BE.** A Western Blot was performed using an anti-TrwC antibody. TrwC, TrwC-BE and BE correspond with D1210 lysates carrying pTrwC, pTrwC-BE and pBE respectively, after 3 hours of induction with IPTG (for pTrwC) or aTc (for pTrwC-BE and pBE). The expected sizes for the proteins are: TrwC 108kDa, TrwC-BE 290 kDa, BE 183 kDa. Left lane, molecular weight marker. Black arrows point to TrwC and TrwC-BE bands.

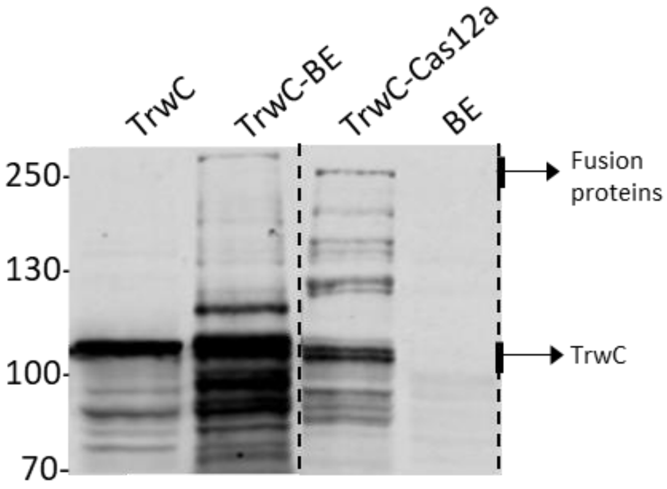
